# Extracellular vimentin as a target against SARS-CoV-2 host cell invasion

**DOI:** 10.1101/2021.01.08.425793

**Authors:** Łukasz Suprewicz, Maxx Swoger, Sarthak Gupta, Ewelina Piktel, Fitzroy J. Byfield, Daniel V. Iwamoto, Danielle Germann, Joanna Reszeć, Natalia Marcińczyk, Robert J. Carroll, Marzena Lenart, Krzysztof Pyrc, Paul Janmey, J.M. Schwarz, Robert Bucki, Alison Patteson

## Abstract

Infection of human cells by pathogens, including SARS-CoV-2, typically proceeds by cell surface binding to a crucial receptor. In the case of SARS-CoV-2, angiotensin-converting enzyme 2 (ACE2) has been identified as a necessary receptor, but not all ACE2-expressing cells are equally infected, suggesting that other extracellular factors are involved in host cell invasion by SARS-CoV-2. Vimentin is an intermediate filament protein that is increasingly recognized as being present on the extracellular surface of a subset of cell types, where it can bind to and facilitate pathogens’ cellular uptake. Here, we present evidence that extracellular vimentin might act as a critical component of the SARS-CoV-2 spike protein-ACE2 complex in mediating SARS-CoV-2 cell entry. We demonstrate direct binding between vimentin and SARS-CoV-2 pseudovirus coated with the SARS-CoV-2 spike protein and show that antibodies against vimentin block *in vitro* SARS-CoV-2 pseudovirus infection of ACE2-expressing cells. Our results suggest new therapeutic strategies for preventing and slowing SARS-CoV-2 infection, focusing on targeting cell host surface vimentin.

## I. Introduction

Infection of human cells by pathogens, including SARS-CoV-2, proceeds by a series of cell surface protein binding and membrane fusion events that are usually centered on a crucial receptor. The SARS-CoV-2 virus is genetically similar to SARS-CoV (SARS) and uses the SARS-CoV receptor, angiotensin-converting enzyme 2 (ACE2), for cell entry (*1, 2*). The ACE2 receptor is expressed in a plethora of tissues, including the lung, kidney, gastrointestinal tract, and vascular endothelium, which all serve as sites for SARS-CoV-2 infection(*3*). While ACE2 seems to be required for SARS-CoV and SARS-CoV-2 infection, it does not appear solely sufficient. The expression of ACE2 in the human respiratory system is low compared to other organs (*4–6*) and while the affinity of the SARS-CoV2 spike protein with ACE is especially strong, the binding-on rate is slow (*1, 7*). At the super-physiological concentrations above nM used *in vitro*, the half time of maximal binding for SARS-CoV-2 is around 30 s, and the concentration in vivo is substantially lower. These findings have given rise to an emerging hypothesis of critical co-receptor that facilitate binding of the SARS-CoV virus and its delivery to ACE2 (*8*), and several possible SARS-CoV-2 co-receptors candidates have been found, including neuropilins (*9*), heparan sulfate (*10*), and sialic acids (*11*). The ongoing COVID-19 pandemic and the threat of future coronavirus outbreaks underscore the urgent need to identify the precise entry mechanism used by the SARS-CoV-2 virus to develop protective strategies against them.

Here, we report that cell surface vimentin acts as a critical co-receptor for SARS-CoV-2 host cell invasion and that antibodies against vimentin can block up to 80% of the cellular uptake of SARS-CoV-2 pseudovirus. While cell surface vimentin is an unconventional target for viruses, there are now numerous studies implicating its role in the binding and uptake of multiple different viruses (*12–19*), including the SARS-CoV virus (*20*), suggesting it might also be involved in cell host invasion by SARS-CoV-2. Interestingly, the expression of SARS-CoV-2 entry factors, ACE2 and TMPRSS2, is particularly high in nasal epithelial goblet secretory cells and ciliated cells (*21, 22*), on which immunohistological studies have shown the presence of vimentin (*23*). We show here that extracellular vimentin is also present in healthy adult lung tissue and detail the numerous routes by which it might arise in the lung, the respiratory track, and other tissues. We demonstrate that vimentin binds to SARS-CoV-2 pseudoviruses that are equipped with SARS-CoV-2 spike 2 protein via dynamic light scattering and atomic force microscopy and propose a novel mechanism in which non-vimentin expressing cells can acquire vimentin released into the extracellular environment by neutrophil netosis. Our work critically highlights extracellular vimentin as a potential target against SARS-CoV-2 that could block the spread of COVID-19 and potentially other infectious diseases caused by viruses and bacteria that exploit cell surface vimentin for host invasion.

## II. Results

### Presence of extracellular vimentin in human lung, airway fluids, and fat tissue

Vimentin is an unexpected target for SARS-CoV-2 viral entry into host cells lining the nasal and lung epithelial airways (Figure 1). Intermediate filaments (IFs) are categorized into five types based on similarities in sequence, which also exhibit similarities in tissue origin (*24, 25*). Keratin is the main IF protein expressed in epithelial cells, whereas vimentin is expressed in mesenchymal cells such as fibroblasts, endothelial cells and leukocytes. While vimentin is not nascently expressed in epithelial cells, its expression can occur in transformed cells associated with cancer, fibrosis, or immortalized cell lines.

**Figure 1.**
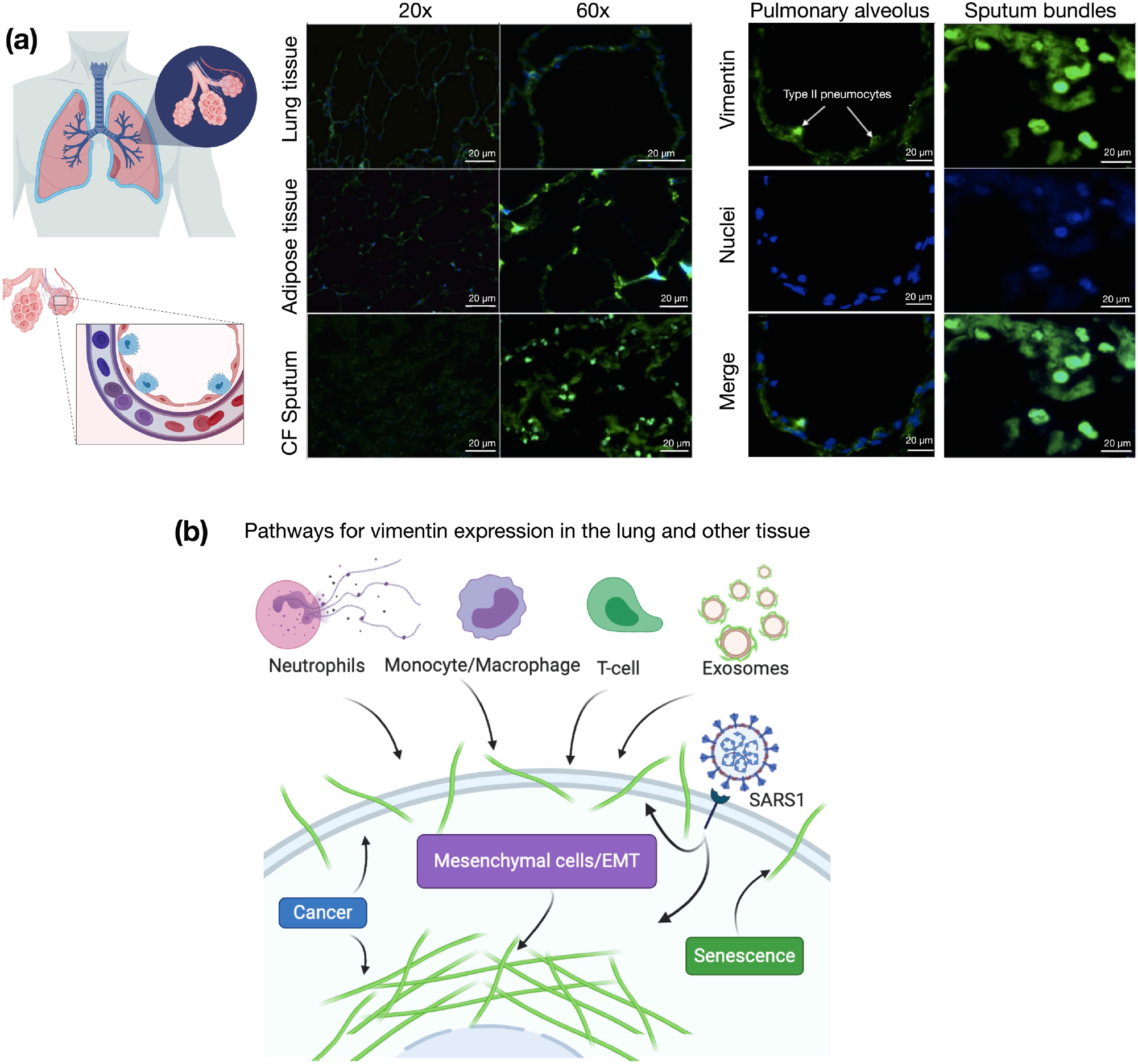
Presence of extracellular vimentin in human lung, airway fluids, and fat tissue. **(a)** Positive staining for extracellular vimentin (green) in human lung, fat tissue, and sputum obtained from cystic fibrosis (CF) patients. Vimentin appears on the apical side of type I and type II pneumocytes. DNA stained with DAPI. **(b)** There are numerous internal and exogenous pathways by which vimentin may be found in lung epithelia and other tissues, in either intracellular or cell surface forms (shown as green filaments). Vimentin is expressed directly by mesenchymal cells, cells having undergone EMT, cancer cells, senescent fibroblasts, and interestingly by cells bound and infected by the SARS-CoV virus (see Table 1). Exogenous sources of vimentin are largely related to immune response and tissue injury in the form of vimentin exported by neutrophils, T-lymphocytes, monocytes/macrophages, and exosomes. Schematics generated with Biorender.com.

There are however many routes by which vimentin might occupy the extracellular space of lung and other tissues. Lung epithelial cells are capable of expressing vimentin. This occurs, for instance, if the cell becomes fibrotic, cancerous, or undergoes the epithelial to mesenchymal transition. However, the source of extracellular vimentin need not be the lung epithelial cell itself, as other cell types have active mechanisms for releasing vimentin into the extracellular space (Fig. 1). This especially includes neutrophils (*26, 27*) macrophages (*28*), and endothelial cells (*29, 30*). While early studies have attributed extracellular vimentin to cytoskeletal debris and disruptions in the plasma membrane, it is now clear that it appears in the absence of cell damage (*31*). While the functions of cell surface vimentin are still largely mysterious, cell surface vimentin has been shown to act as a biochemical signal between different cell types but also serves as an attachment factor through which multiple types of bacteria and viruses infect cells (*32, 33*).

The presence of vimentin on the surface of alveolar pneumonocytes and in the extracellular space of the lungs was assessed with an anti-vimentin antibody by immunostaining healthy lung parenchyma without pathology and sputum samples collected from the respiratory tract of cystic fibrosis patients (Fig. 1a). The presence of vimentin on the apical surface of type I and type II pneumocytes, cells without endogenous vimentin expression, indicate its extracellular origin. Immunostaining of airway sputum from cystic fibrosis patients showed the presence of vimentin in DNA-rich aggregate structures. Since obesity was identified as an important risk factor for complications in patients that suffered from SARS-CoV-2 infection (*34*), we also assessed the presence of vimentin in human adipose tissue (Fig. 1a). Indeed, vimentin expression was present in the stromal tissue, not in the adipocytes. The identification of extracellular vimentin in lung, sputum, and adipose tissue indicates a surprising pattern of extracellular vimentin presence in interstitial tissue and its possible role in SARS-CoV-2 uptake in lung and other ACE2 expressing tissues.

### Binding of vimentin to the SARS-CoV-2 spike protein

Next, we tested whether vimentin could bind to SARS-CoV-2 pseudoviruses displaying the antigenically correct spike protein in vitro. These pseudovirus present the SARS-CoV-2 spike protein on replication-incompetent virus particles that contain a heterologous lentiviral (HIV) core (Integral Molecular, RVP-701G). Binding of purified bacterially-expressed human vimentin to these pseudoviruses was measured by a combination of dynamic light scattering (DLS) and imaging by atomic force microscopy (AFM). Figure 2A shows that the hydrodynamic radius of the pseudovirus was 60 nm, consistent with the expected size of a lentivirus similar to the size of SARS-CoV-2 (*47*). As purified vimentin was added to the pseudovirus suspension, its hydrodynamic radius increases to approximately 150 nm. Here, the increase in apparent size is presumably due the bridging together of pseudovirus by vimentin oligomers. We estimate that at the highest concentration of vimentin there is an excess of 6 x 10^4^x vimentin oligomers to spike protein (Methods). The increase in apparent size is not due to a separate contribution of large vimentin filaments to the mixture, because scattering from vimentin alone was negligible at all concentrations compared to that of the pseudovirus, and the increase in scattering of the mixtures is larger than the sum of separate contributions from pseudovirus and vimentin (SI Fig. 1). Separate measurement of the vimentin used for these studies showed that its effective hydrodynamic radius was 100 nm, which represents small oligomer of vimentin similar in size to those reported at the cell surface (*48*).

**Figure 2.**
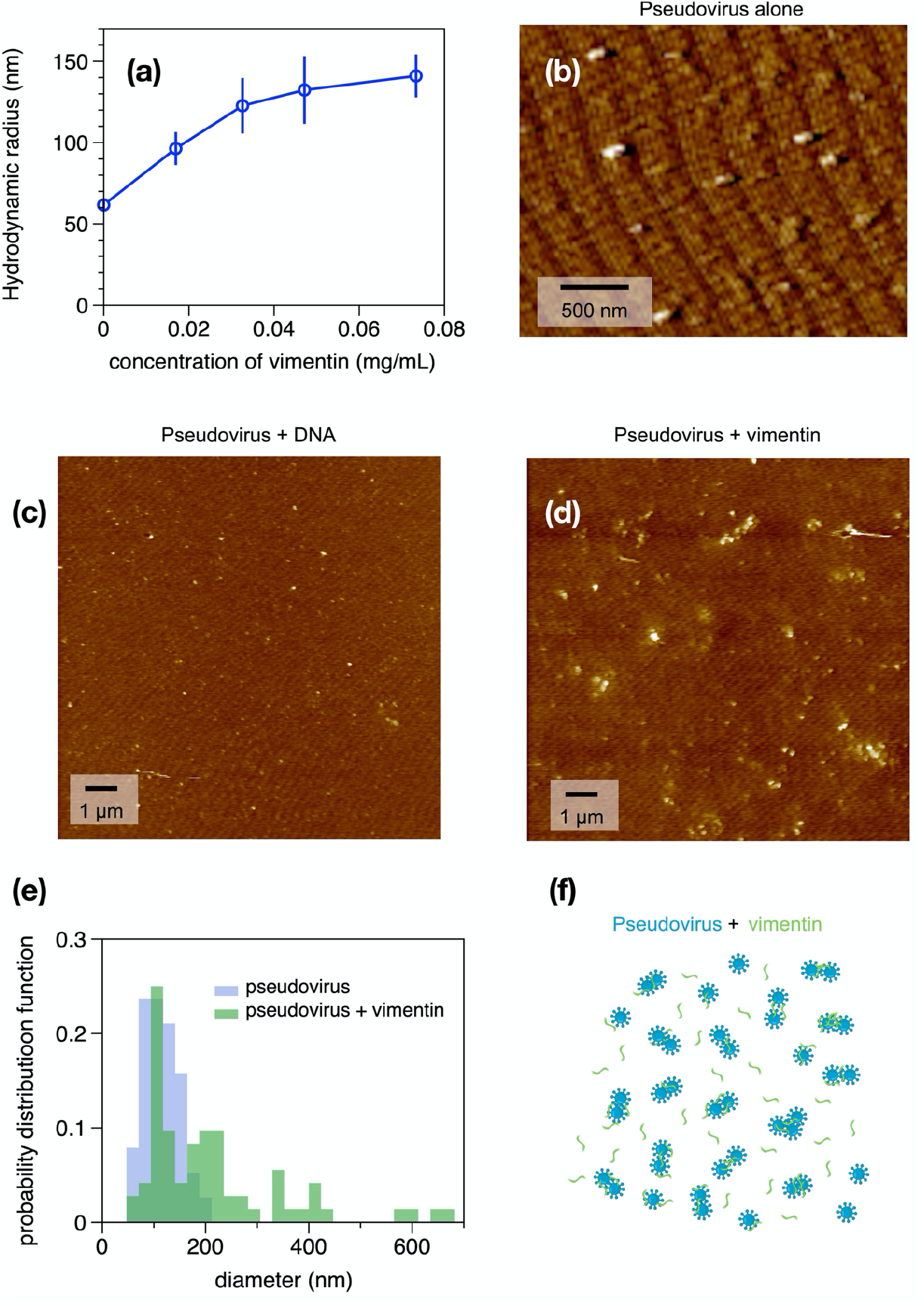
Binding of vimentin to SARS-CoV-2 pseudovirus. Purified human recombinant vimentin was added to suspensions of SARS-CoV-2 spike protein-containing pseudovirus, and their size was measured by dynamic light scattering (a) and atomic force microscopy (b-e). The size of the pseudovirus increased from 60 nm to 150 nm after addition of 0.07 mg/ml vimentin (panel a). Panel b-d: The pseudoviruses were imaged using atomic force microscopy before and after addition of either DNA (0.08 mg/mL) or vimentin (0.07 mg/ml). The probability distribution functions (e) show that the average size of pseudovirus imaged by AFM confirms the change in size detected by DLS. (f) Schematic representation indicating how binding of vimentin to SARS-CoV-2 spike protein could couple particles together and increase their effective radii.

The binding of vimentin to the pseudoviruses is likely specific and not a result of the highly-charged polyelectrolyte nature of vimentin. This was tested by using double-stranded DNA, a biopolymer of similar size and surface charge, which can also be found in the extracellular space of tissue. Addition of double stranded DNA did not lead to an increase pseudovirus size (SI Fig. 2).

The samples used for DLS were also examined by atomic force microscopy. Figure 2 shows the pseudovirus alone (Fig 2b) and pseudovirus after addition of DNA (Fig. 2c) or vimentin (Fig. 2d). Histograms of the size distributions (Figures 2e and f) show pseudovirus diameters consistent with the radii measured by DLS and that after addition of vimentin, some single pseudoviruses appear larger, but small clusters of pseudoviruses, possibly bridged by vimentin oligomers, are common (Fig. 2f). Binding of vimentin oligomers or short filaments to the pseudovirus was selective for vimentin; we found that similar increases in pseudovirus size after titration with three different preparations of vimentin, but no significant increase after addition of double stranded DNA, a polymer that is also found extracellularly with similar size and surface charge of vimentin (SI Fig. 3). Overall, the data in Figure 2 shows that purified vimentin can bind the SARS-CoV-2 spike protein-containing pseudoviruses and cause it to aggregate.

### Anti-vimentin antibodies block uptake of SARS-COV-2 pseudoviruses in cultured human epithelial cell lines

If extracellular vimentin can bind to SARS-CoV-2 virus and help capture it at the cell surface, then vimentin might increase SARS-CoV-2 host cell invasion by acting as an attachment factor, and anti-vimentin antibodies could block its uptake. To test this idea, we identified two model cell lines, human embryonic kidney epithelial cells HEK 293T-hsACE2 (Figure 3) and human lung cancer cells A549-hsACE2 (Figure 4), both stably expressing the ACE2 receptor to enable host cell invasion by the SARS-CoV-2 virus or pseudoviruses. Both of these epithelial cell lines express vimentin, though the presence of cell surface vimentin was unclear. Thus, we performed immunofluorescence studies comparing cells treated with primary anti-vimentin antibodies before fixation versus after fixation and cell membrane permeabilization by Triton X-100 (Methods). We compared our results with mouse embryonic fibroblasts that form intracellular filamentous vimentin networks, emblematic of mesenchymal cells (Fig. 4a). In both the HEK 293T-hsACE2 and A549-hsACE2, the presence of cell surface vimentin was detected.

**Figure 3.**
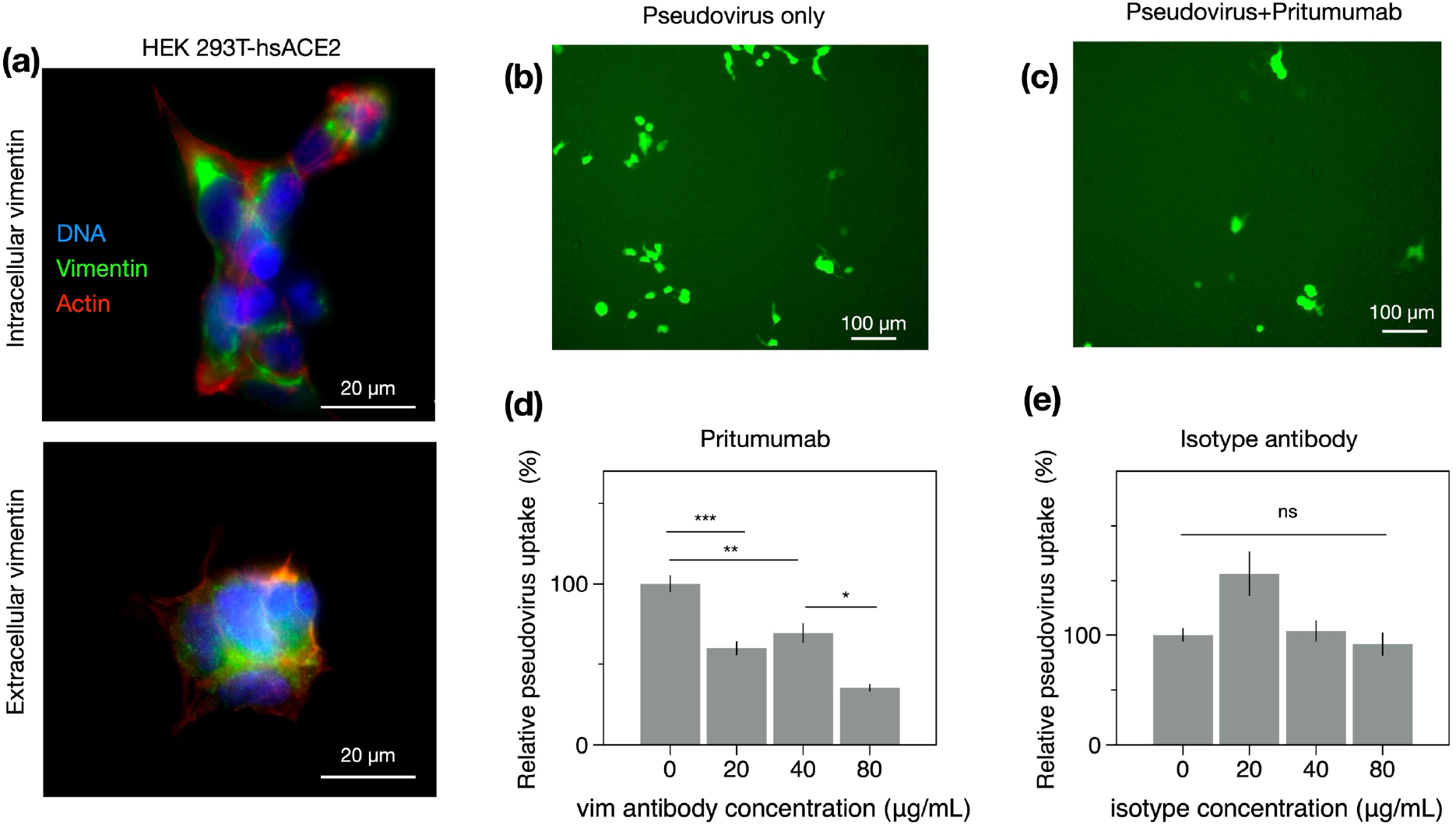
Anti-vimentin antibodies block uptake of SARS-CoV-2 pseudovirus in HEK 293T-hsACE2. **(a)** Positive staining for extracellular vimentin in mouse embryonic fibroblasts and human kidney epithelial cells HEK 293T-hs ACE2 and mouse embryonic fibroblasts. Images show vimentin (green), actin (red), and DNA (blue). To stain for extracellular vimentin, cells were exposed to a primary polyclonal chicken anti-vimentin antibody, before fixation, then permeabilized and stained for actin. (**b)** Schematic of experimental design of pseudovirus infection studies. HEK cells are pre-exposed to the anti-vimentin antibody Pritumumab before infection by pseudovirus bearing the SARS-CoV-2 spike protein and a GFP reporter. **(c)** Representative fluorescence images showing cells expressing GFP after pseudovirus exposure with and without Pritumumab treatment. **(d)** Pritumumab inhibits cellular infection by up to 60%. **(e)** Use of an isotype antibody does not inhibit infection, corroborating a specific interaction between the SARS-CoV-2 spike protein and extracellular vimentin. **(f)** Use of another vimentin antibody, a chicken IgY polyclonal antibody, also blocks cellular transduction by SARS-CoV-2 pseudovirus. Denotations: *, P ≤ 0.05; **, P < 0.01; ***, P < 0.001; NS, P > 0.05.

**Figure 4.**
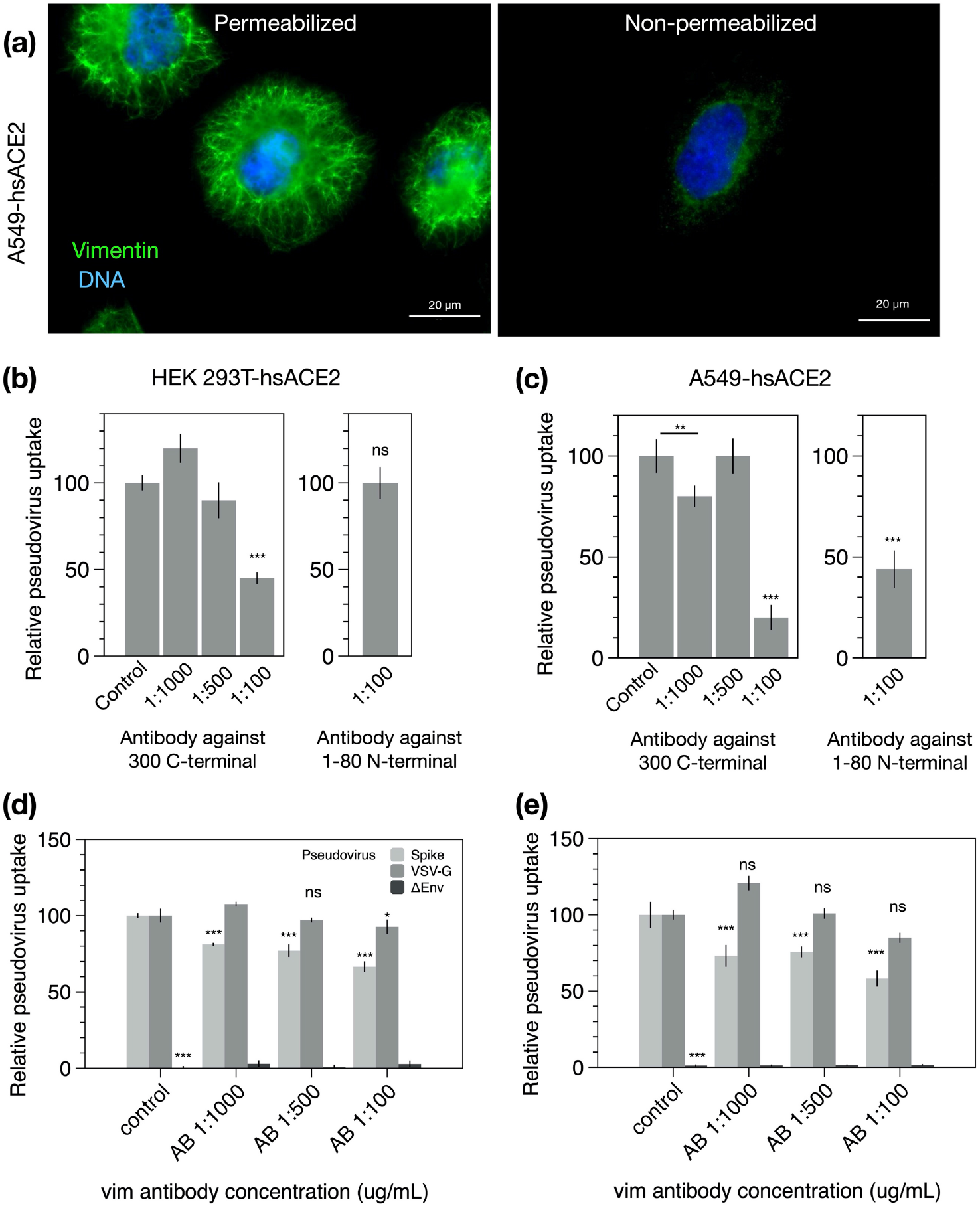
SARS-CoV-2 pseudovirus transduction results based on plate readers for HEK 293T-hsACE2 and A549-hsACE2 cells. **(a)** Extracellular vimentin was detected on A549-hsACE2 as shown by immunofluorescence images. **(b)** The recombinant Anti-Vimentin antibody – rabbit monoclonal IgG, which binds to the vimentin C-terminus blocks uptake in both HEK 293T-hsACE2 and **(c)** A549-hsACE2 cells. The primary anti-vimentin polyclonal rabbit antibody that targets the phosphorylation site of SER56 on the N-terminus blocks uptake in A549-hsACE2 cells but not HEK 293T-hsACE2 cells. Denotations: *, P ≤ 0.05; **, P < 0.01; ***, P < 0.001; NS, P > 0.05.

Next, host cell invasion studies were conducted using SARS-CoV-2 pseudoviruses bearing a green fluorescent protein (GFP) mRNA reporter. Cultured cells were exposed to varying concentrations of pseudoviruses (Methods), and the resulting number of transduced cells was detected by epifluorescence microscopy of cell GFP expression and monitored over the course of three days. As expected, the number of GFP-expressing cells reached a maximum three days after pseudovirus exposure and was pseudovirus load dependent. Maximum infection rates were in the range of 1 to 10% among experimental replicates for HEK 293T-hsACE2 and A549-hs ACE2 cells.

Can anti-vimentin antibodies block SARS-CoV-2 uptake and does inhibition depend on the antibody’s vimentin-binding epitope? To address this question, we presented cells with a number of different antibodies against vimentin (see Table 1) and measured the resultant change in transduction compared to the case without antibodies. We began these studies with the most promising therapeutic anti-vimentin antibody, pritumumab, a human-derived IgG antibody, reported to be specific against extracellular vimentin and known to bind to vimentin’s C-terminal domain (*49*). In this case, we pre-exposed HEK 293T-hsACE2 cells with various concentrations of Pritumumab prior to infection by SARS-CoV-2 pseudoviruses and found a dose-dependent decrease in infection efficiency. We found a maximum reduction of pseudovirus transfection of up to 60% when cells were incubated with 120 μg/ml Pritumumab (Fig. 3d). The specificity of the pritumumab interaction with extracellular vimentin was tested against an IgG isotype antibody control (Fig. 3e), which did not block entry of the SARS-CoV-2 pseudoviruses, suggesting a specific interaction between vimentin and the SARS-CoV-2 spike protein.

**Table 1.** Sources of intracellular and exported vimentin.

| Source | Description | Reference |
| --- | --- | --- |
| ***Extracellular Sources – Largely implicated in immune and tissue injury*** |  |  |
| Neutrophils | Released during apoptosis or NET formation; Source of serum citrullinated vimentin commonly found in rheumatoid arthritis | (27, 35) |
| Monocytes/Macrophages | Secreted after simulation by TNF-α or oxidized low density lipoprotein; possibly tissue injury; phosphorylated | (28, 36) |
| T-lymphocytes | Released on apoptosis; Binds phospholipase II to promote arachidonic acid metabolism | (37) |
| Exosomes | Astrocytes, activated by injury, can produce extracellular vimentin in the form of exosomes that are delivered to neurons; pre-adipocytes | (38-40) |
| ***Sources from the cell itself – Mesenchymal and transformed cells*** |  |  |
| Mesenchymal cells/EMT | Vimentin is expressed endogenously in mesenchymal cells and is a wide-spread marker EMT | (41, 42) |
| Cancer | Intracellular: Upregulation of vimentin in multiple types of cancer; Extracellular: Target for isolating circulating tumor cells, designing vaccines and enhanced chemotherapy; sometimes proteolyzed | (43, 44) |
| Senescent Fibroblasts | Post-translational modification of cysteine 328 by malondialdehyde in senescent cells leads to its secretion and surface exposure | (45) |
| SARS-CoV Stimulation | Extracellular vimentin expression increases upon SARS-CoV binding to cell surface; SARS-CoV also upregulates cytoplasmic vimentin by the TGF-β pathway to promote a fibrotic response in lung cells | (20, 46) |

Next, we tested the efficiency of other anti-vimentin antibodies (Table 2) in blocking host cell invasion by SARS-CoV-2 pseudoviruses and sought a vimentin binding domain that could be targeted against SARS-CoV-2 (Table 2). We used: (i) a chicken polyclonal antibody that binds to multiple epitopes; (ii) anti-vimentin antibodies targeting parts of the C terminus: in addition to Pritumumab, a human vimentin IgG against amino acid 300; and (iii) antibodies against the N-terminus: a rabbit monoclonal antibody corresponding to residues surrounding Arg45 and a polyclonal rabbit antibody IgG around the non-phosphorylation site of Ser56.

**Table 2.** List of anti-vimentin antibodies.

| Anti-Vimentin antibody | Binding domain on vimentin | Effectiveness in blocking uptake of SARS-CoV-2 pseudovirus |
| --- | --- | --- |
| Pritumumab (Nascent Biotech); A natural human IgG1 kappa antibody was obtained from a regional draining lymph node of a patient with cervical carcinoma | Epitope is the C2 core region (coil 2 of the central rod) of cell-surface vimentin (49, 50) | Blocks uptake in HEK 293t-hsACE2 up to 60% |
| Chicken polyclonal IgY antibody (Novus Biologicals)- Cat# NB300-223 | Polyclonal antibody, multiple epitopes | Blocks uptake in HEK 293t-hsACE2 up to 80% |
| Rabbit Monoclonal (Cell signaling Technology) Cat# #5471 | N-terminal; produced by immunizing with a synthetic peptide corresponding to residues surrounding Arg45 of human vimentin | Does not block uptake in HEK 293t-hsACE2 |
| Recombinant Anti-Vimentin antibody – IgG MW~150kDa [EPR3776] (Abcam) Cat# ab92547 | C-terminal; produced by immunizing with a synthetic peptide containing 17 residues from within amino acids 425-466 of human vimentin | Blocks uptake in HEK 293T-hsACE2 by 60%; A549-hsACE2 up to 80% |
| Primary anti-vimentin polyclonal rabbit antibody IgG ~150kDa (antibodies-online.com) Cat# ABIN6280132 | N-terminal; Synthesized peptide derived from human Vimentin around the non-phosphorylated site of Ser56, Binding Specificity AA 1-80, Ser56 | Blocks up to 50% in A549-ACE2; Does not block in HEK 293T hsACE2 |

We note that some but not all anti-vimentin antibodies blocked SARS-CoV-2 pseudoviruses cellular uptake in a dose-dependent manner. Using epifluorescence microscopy to quantify the pseudoviral transduction in HEK 293T-hsACE cells, we found that the polyclonal antibody against vimentin decreased host cell invasion by up to 80% (Fig. 3f), whereas the rabbit monoclonal antibody targeting the N-terminal was not effective at blocking uptake (SI Fig. 4). As an independent measurement, we used plate readers to measure GFP expression due to SARS-CoV-2 pseudovirus transfection in both HEK 293T-hsACE and A549-hsACE cells (Fig. 4). In this case, the recombinant antibody against the C-terminal 41-amino acids and the polyclonal rabbit antibody targeting Ser56 of the vimentin N-terminal were used. The C-terminal antibody was efficient in blocking uptake up 60% in HEK 293T-hsACE and up to 80% in A549-hsACE. The results on the Ser56 antibody were somewhat mixed: this antibody did not block uptake in HEK 293T-hsACE2 but did block up to 55% in A549-hsACE2. Taken together, our results suggest that antibodies targeting the C-terminal tail domain of vimentin are most efficient in blocking SARS-CoV-2 host cell invasion. Antibodies against the N-terminus are generally not as efficient but may work to block uptake in some cases (e.g. A549-hsACE2 in Fig. 4b).

To further test the specificity of vimentin with the SARS-CoV-2 spike protein (rather than the pseudovirus in general), we performed a control experiment comparing pseudoviruses harboring SARS-CoV-2 proteins with those carrying VSV-G (positive control, no spike protein) as well as with pseudoviruses lacking the fusion protein (ΔEnv – negative control) (Table 3). Each pseudovirus was bearing plasmid encoding GFP. A549-hsACE2 and HEK293T-hsACE2 cells were incubated with varied dilution of rabbit monoclonal antibody specific to human vimentin C-terminal amino acids 425-466.

**Table 3.** List of pseudoviruses.

| Type of pseudovirus | Mechanism of cell entry | Interaction with vimentin |
| --- | --- | --- |
| Spike-GFP – lentiviral packaging plasmid (psPAX) with SARS-CoV-2 S glycoprotein (pCAGGS-SARS-CoV-2-S) and third plasmid encoding GFP protein (Lego-G2)<br>Spike = psPAX + Lego-G2 + pCAGGS-SARS-Cov-2-S | SARS-CoV-2 spike glycoprotein directly binds with the host cell surface ACE2 receptor and TMPRSS2 (51), possible direct attachment with surface vimentin acting as co-receptor | Anti-vimentin antibodies (17 C-terminal residues aa 417-426) inhibited Spike-GFP pseudovirus entry into A549hsACE2 cells by 35% and HEK 293ThsACE2 cells by 45% in dose-dependent manner |
| SARS-CoV-2 Wuhan-Hu-1 strain (#RVP-701G, Integral Molecular) – heterologous lentiviral core, three separate plasmids encoding lentiviral gag protein, spike protein and reporter gene |  | Pritumumab blocked up to 60% of viral transduction in kidney epithelial cells expressing ACE2. High inhibition efficacy of anti-vimentin antibodies to C-terminal, developed both in rabbit and chicken. |
| VSV-GFP – lentiviral packaging plasmid (psPAX) with the VSV-G envelope plasmid (pMD2G) and plasmid encoding GFP VSV-G = psPAX + Lego-G2 + pMD2G | VSV bind to low-density lipoprotein receptor (LDL-R) to entry the cells, considered as positive control (52) | Vimentin antibodies do not block VSV-GFP pseudovirus uptake in A549-ACE2 nor HEK 293ThsACE2 cells |
| ΔEnv-GFP – lentiviral plasmid (psPAX) with green fluorescent protein (GFP) encoding plasmid (Δenv = psPAX + Lego-G2) | Lack of envelope protein, do not enter into the cells, considered as negative control | No direct or indirect interactions with vimentin recorded |

Inhibition of uptake of S-SARS-CoV-2 pseudovirus was recorded at up to 35% and 75% for lung-derived and kidney-derived epithelial cells, respectively. Moreover, C-terminal antibody inhibition exhibit dose-dependent manner in both cell lines. In case of VSV-G pseudovirus, negligible inhibition was noted what indicates specificity of anti-vimentin antibody effect to inhibit cell surface vimentin-mediated SARS-CoV-2 cell entry and its potential interaction with glycoprotein S. Negative control (ΔEnv) did not differ from background measurement consisted of cells in cell culture medium without pseudoviruses.

### Cell surface vimentin acquisition from the extracellular environment

Our results thus far suggest that extracellular vimentin might bind to the SARS-CoV-2 spike protein and enhances host cell invasion by the SARS-CoV-2 virus. Studies on SARS-CoV and SARS-CoV-2 indicate that while ACE2 is required for infection, it is not solely sufficient for cellular uptake and invasion (*8*). We therefore posit that extracellular vimentin is a key player in SARS-CoV-2 invasion and that its presence in the lung and other tissues is an important pre-condition of the extracellular environment that enhances SARS-CoV-2 infection. As a pre-requisite, we have identified that vimentin is present in extracellular space of healthy adult lung and fat tissue (Fig. 1). Vimentin may appear on the cell surface and in the extracellular space of the lung via numerous routes (Fig. 1, Table 1). These routes are likely associated with inflammatory response in the course of disease (*e.g*. fibrosis and cancer) but might also reflect homeostatic regulation associated with normal tissue functions and maintenance. In the context of the lung, the most probable sources are vimentin-expressing lymphocytes, macrophages and neutrophils that specialize in fighting off pathogens.

Here, we demonstrate a novel mechanism by which non-vimentin expressing cells acquire cell surface vimentin from the extracellular environment via neutrophil NETosis (*26, 35*). NETosis represents one of the mechanisms of host defense, in which neutrophils expel large webs of DNA that entrap bacteria. Additionally, during NETosis disassembly of the vimentin network take place. NETosis requires activation of peptidyl arginine deiminase 4 (PAD4), which citrullinates histones in the process of releasing DNA from neutrophil heterochromatin. At the same time, the cytoskeleton disassembles to allow the nuclear content released after nuclear membrane rupture to reach the plasma membrane and eventually enter the extracellular space as the plasma membrane ruptures (*35*). Disassembly of the vimentin network is also promoted by citrullination, and vimentin is a major substrate for PADs (*26*). Citrullination leads to disassembly of the vimentin network, which can then be released into the extracellular space and become accessible to the cell surfaces of other tissues.

To show that neutrophil-released vimentin can be acquired on the surface of other cell types, we stimulated NETosis in neutrophils with phorbol myristate acetate (PMA), collected the supernatant following centrifugation, and then incubated the supernatant with a different cell type (Fig. 5a). Here we used vimentin-null mouse embryo fibroblasts (mEF), so that any vimentin detected on their surface was indicative of vimentin acquired from the extracellular environments. As shown in Fig. 5b, after exposing cells to the supernatant containing NETs (positive for DNA and elastase; data not shown), the vimentin-null cells stained positive for cell-surface vimentin in immunofluorescence microscopy. These results demonstrate a new mechanism by which vimentin is released into the extracellular environment of tissues, where it can then bind to attach the extracellular surface of cells. Treatment of cells with DNASE did not prevent vimentin deposition on cells (SI. Fig. 6), indicating that vimentin was not binding to the cell surface through DNA, which is released upon NETosis. Nor did treatment of cells with hyaluronidase, which degrades the cell glycocalyx, release cell surface vimentin (SI. Fig. 6).

**Figure 5.**
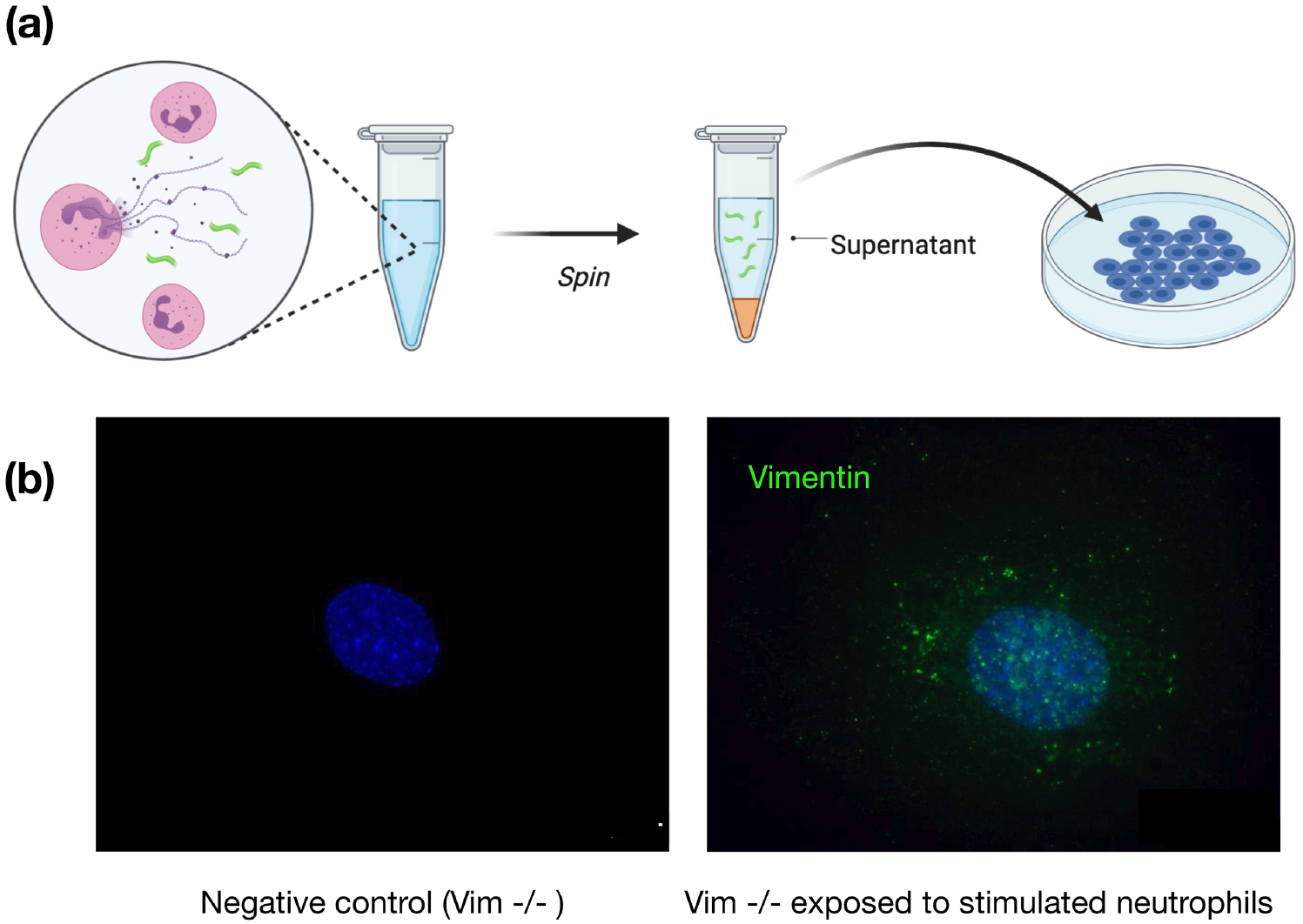
Acquisition of cell surface vimentin from extracellular environment. **(a)** Neutrophil NETosis stimulates disassembly of the vimentin network and enables its release into the extracellular space. **(b)** Immunofluorescence images of vimentin-null mEF staining positive for extracellular vimentin after exposure to supernatant of NETosis-activated neutrophils, indicating the acquisition of extracellular vimentin by cells that do not express vimentin. Schematics generated with Biorender.com.

### Modeling the interactions between cell surface vimentin and SARS-CoV-2

Both SARS-CoV and SARS-CoV-2 can enter host cells via both membrane fusion and endocytic pathways; although, both pathways are dependent upon ACE2 binding (*51, 53–57*). Emerging evidence indicates that membrane fusion is the dominate pathway when the transmembrane protease serine 2 TMPRSS2 is present on the cell surface (*58*). However, in the absence of such proteases, coronaviruses will use clathrin- and non-clathrin-mediated endocytosis (*54, 59*). TMPRSS is expressed in lung cells (*60, 61*), but during later stages of SARS-CoV-2 infection the virus spreads to different tissues and infects other cell types (*62, 63*). If extracellular vimentin traps the SARS-CoV-2 virus at the cell membrane, it might alter the dynamics of virus at the cell membrane and even mediate membrane fusion or endocytic events in different cell types.

Here, we developed a computational model to simulate the interactions between SARS-CoV-2 virus, extracellular vimentin, and cell membrane surface (Fig. 6). The virus is modeled as sphere with multiple spikes. We choose 200 spikes as an upper estimate on the number of spike proteins observed in SARS-CoV-2 (*64*). The cell membrane is modeled as a thermally-fluctuating elastic sheet, which is either purely elastic with no bending rigidity (0 k_B_T) or elastic with finite membrane rigidity (40 k_B_T). Cell surface vimentin is modeled as a set of 40 nm semiflexible polymers, based on recent work showing cell surface vimentin is mainly non-filamentous and in the form of oligomers (4-12 monomers and tetramers) (*48*). The cell surface vimentin is bound to the cell membrane and can bind to the spike protein. The model begins with the virus attached to the cell membrane via an ACE-2 binding site. For modeling details, see Methods and SI materials.

**Figure 6.**
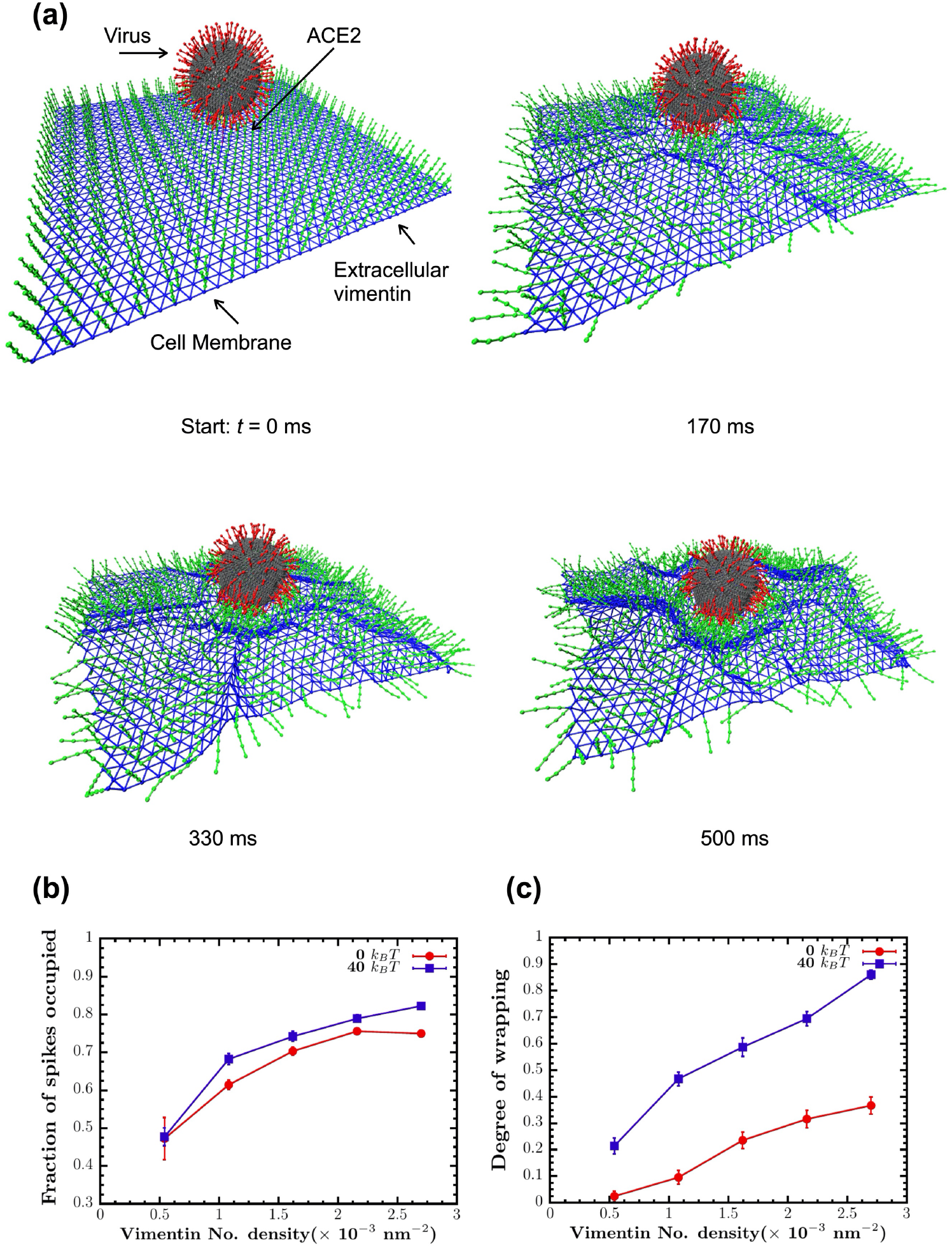
Vimentin’s involvement in spike 2-ACE2 interactions. **(a)** Cell surface vimentin acts as a co-receptor that enhances binding to the SARS-CoV-2 virus in either direct fusion or endocytic pathways to enhance wrapping and endocytosis of the virus. A molecular dynamics simulation of the SARS-CoV-2 virus at an elastic shell membrane shows that binding of extracellular vimentin with the virus spike protein facilitates wrapping of cell membrane around the virus. Both the fraction of spikes bound to surface vimentin **(b)** and the degree of membrane wrapping **(c)** increases as the number density of surface vimentin increases. Finite membrane bending rigidity (40 k_B_T, blue squares) enhances wrapping compared to the case without bending rigidity (0 k_B_T, red circles).

As shown in Fig. 6, we find that these simple interaction rules drive the cell membrane to partially wrap around the virus and in effect allow the virus to interact with an increased surface area of the cell membrane. Increasing cell surface vimentin increases the fraction of spikes bound to cell surface vimentin and increases the effective degree of wrapping around the cell virus (Methods). Interestingly, finite cell membrane bending rigidity enhances wrapping of the virus, as compared to the case where the cell membrane is purely elastic. These results presented in Fig. 6 indicate that co-receptors, such as cell surface vimentin, increases the effective interactions between the SARS-CoV-2 virus and the cell membrane. In acting as an attachment factor, cell surface vimentin would enhance the probability of a viral fusion event and could perhaps stimulate cell membrane wrapping around the virus (Fig. 6), initiating the first steps towards viral endocytosis.

## III. Discussion

Identifying targets against SARS-CoV-2 is critical because evidence suggests that viral load is correlated to the severity of COVID-19 infection in patients (*65*). Here, we have shown that extracellular vimentin promotes SARS-CoV-2 uptake in lung and kidney epithelial cells and antibodies targeted against vimentin block it. Thus, we propose that vimentin-blocking antibodies could serve as a promising therapeutic strategy against SARS-CoV-2 and future coronaviruses, decreasing viral load, and subsequent patient symptoms.

Although generally considered primarily an intracellular, cytoskeleton protein, a large body of evidence shows that vimentin can also be found on the external surface of cells and in extracellular fluids. Numerous pathways have been identified for controlled release of vimentin often by inflammatory cells. Extracellular release depends on covalent modifications of vimentin, often by phosphorylation or citrullination (*31*). These modifications disassemble cytoskeletal vimentin intermediate filaments and lead to release of vimentin either on the external surface of the same cell or in soluble form or on the surface of exosomes where it can target cells such as epithelial cells (*66*) or neurons (*39, 40*) that do not express vimentin endogenously. Cell surface vimentin is reported to function normally during some types of wound healing (*40, 66, 67*) but is also often increased during pathological states such as cancer (*68, 69*). Numerous pathogens, both viruses and bacteria, exploit cell surface vimentin to gain entry to the host cell (*12–20*). In bacteria, engagement of vimentin is often due to binding of essential virulence factors (*70*). For viruses that are affected by vimentin, binding depends on coat proteins, and in case of SARS-CoV, the potential binding molecules for vimentin is its spike protein, which is similar to the spike proteins of SARS-CoV-2 (*20*).

The utility of an initial binding event at the cell surface prior to engagement of the viral spike protein by its receptor ACE2 is supported by the binding kinetics of the spike protein from SARS-CoV or SARS-CoV-2 to an immobilized chimera of the ACE2 extracellular domain and the Fc fragment of IgG. Surface plasmon resonance measurements show that although the affinity of the SARS-CoV-2 spike protein is greater than that of SARS-CoV spike protein, the rate of binding is much slower (*1, 7*). Therefore, engagement of the virus by an ACE2 bearing cell would be highly enhanced if the virus was first immobilized by a faster binding event. While the kinetic binding rates between vimentin and the SARS-CoV-2 spike protein have yet to be measured, it is likely much faster than the slow binding time between SARS-CoV-2 spike protein and the ACE2 receptor. Thus, extracellular vimentin would serve as a critical co-receptor, binding and engaging with the SARS-CoV-2 virus at the surface of the cell membrane, enhancing its delivery to the ACE2 receptor. We propose that anti-vimentin antibodies can block this interaction, as shown schematically in Fig. 7.

**Figure 7.**
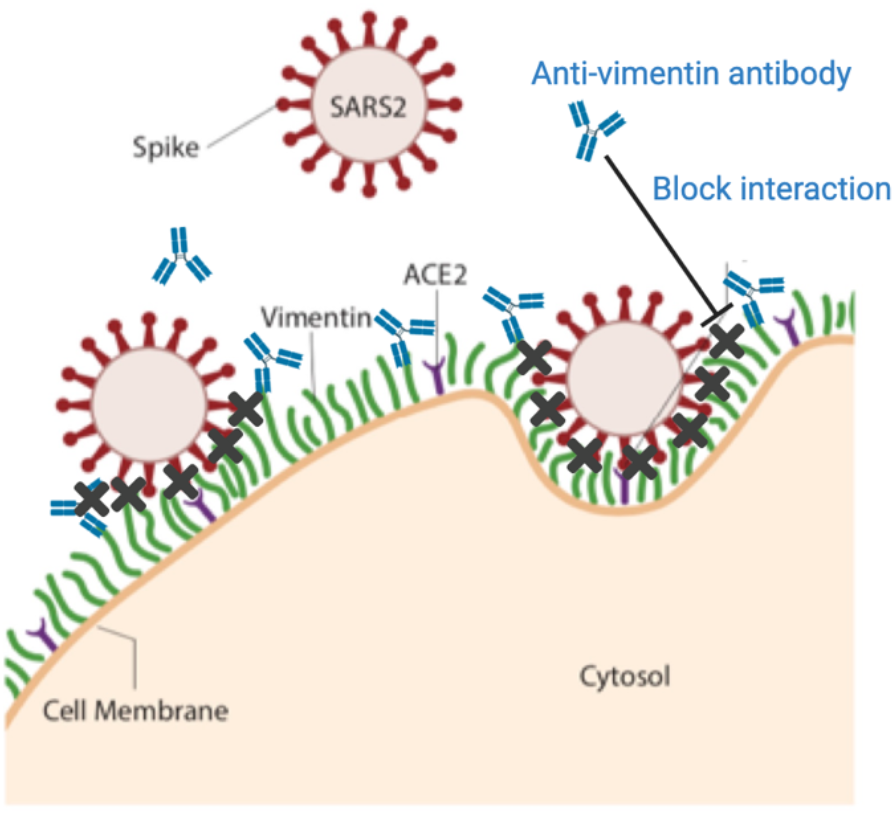
Extracellular vimentin as a potential target for inhibiting SARS-CoV-2 entry. A diagram of proposed mechanism of action. Here, cell surface vimentin (green) acts as a co-receptor that binds to SARS-CoV-2 spike protein. Blocking this interaction via the anti-vimentin antibody Pritumumab reduces cell surface binding of the virus and cellular infection. Schematics generated with Biorender.com.

There are two main pathways by which the SARS-CoV-2 virus is thought to attach and bind to cells for infection. The first is through the transmission of respiratory droplets that enter and move through the nasal and lung epithelial airways. The second way occurs after initial infection by entering into the bloodstream, where it can enter endothelial cells and lead to inflammation and damage in organs such as the heart, kidney, and liver. Thus, our results suggest two general therapeutic strategies to prevent and reduce SARS-CoV-2 infections. The first is to aerosolize the antibody and use it as a nasal spray as a preventive measure against SARS-CoV-2 infection. The second is intravenously post-infection – alone or in combination with other viral inhibitors - to sequester SARS-CoV-2 in the blood and reduce the risk of further tissues beyond the lung getting infected and damaged by the virus.

Finally, while these studies focus on the critical first few steps by which SARS-CoV-2 infects cells, there are *in vitro* and *in vivo* studies that suggest an active role of extracellular vimentin in the cellular response post-infection by SARS-CoV and SARS-CoV-2. For instance, a recent study using Vero E6 cells showed that cell surface vimentin levels increased over 15% after exposure to SARS-CoV pseudovirus (*20*). Another study of over 80 patients with respiratory failure secondary to severe SARS-CoV-2 pneumonia found extracellular vimentin in the bronchoalveolar lavage (BAL) fluid of infected patients (*71*). Vimentin appears to have a complicated role in pathogenesis, acting as both a target for infection (*20, 72, 73*) and an important actor for proper immune response (*26, 28, 74*). Our results here suggest that extracellular vimentin acts as a novel target for SARS-CoV-2 entry inhibition. Identifying the mechanisms that lead to the expression, trafficking, and secretion of vimentin will be pivotal to mediating its role in tissue damage and repair during viral infections.

## IV. Conclusions

Vimentin is secreted into the extracellular environment by multiple cell types and its presence can provide positive signals for wound healing and act as a cofactor for pathogen infection. Both monoclonal anti-vimentin antibodies and a vimentin-specific DNA aptamer developed to isolate circulating cancer cells (*75*) are being applied to infection by SARS-CoV (*20*) and other viruses (*76*). Here, we have demonstrated a new role of extracellular vimentin in SARS-CoV-2 host cell invasion. Further, we find that antibodies against vimentin can block up to 80% of host cell invasion *in vitro*. Our results suggest a new target and therapeutic strategy to reduce SARS-CoV-2 attachment and entry into the cell, which could reduce the spread of the SARS-CoV-2 virus and other possible pathogens that exploit extracellular vimentin.

## V. Methods

### Cell culture

In this study five cell lines were used: wild-type mouse embryonic fibroblasts (mEF vim +/+), vimentin-null fibroblast (mEF vim -/-), adenocarcinomic human alveolar basal epithelial cells transfected with hACE2 (A549-ACE2), provide by Krzysztof Pyrc from Małopolska Centre of Biotechnology; Jagiellonian University; Kraków, Poland) and human kidney epithelial cells HEK293T ACE2-transfected (Integral Molecular, Integral Cat# C-HA101). Cells were cultured in DMEM (ATCC) with 10% of fetal bovine serum (FBS), penicillin (50 μg/mL) and streptomycin (50 μg/mL). Kidney derived cells were additionally treated with 0.5 μg/mL puromycin to maintain ACE2 expression. Cells were maintained at 37°C in an atmosphere containing 5% CO2 with saturated humidity.

### Tissue staining

To evaluate presence of extracellular vimentin on the surface of cells within human tissue, fluorescent immunohistochemistry staining was performed. To do so, paraffin-embedded human lung and fat tissue, as well as sputum obtained from cystic fibrosis patients were cut into 15 μm thick slices on a rotary microtome and placed on glass slides. After overnight drying, slides were rehydrated by immersion in xylene and ethanol for 10 min at each step. Slides were hydrated with PBS and transferred to EDTA-containing antigen retrieval solution (Sigma) for 1h at 100°C. Subsequently, samples were blocked in blocking buffer (1% BSA in PBS) for 30 min at RT. Primary rabbit anti-vimentin antibodies to the C-terminus at 1:200 were placed directly on tissues and left for overnight incubation in humidified chamber at 4°C. Afterwards, slides were washed three times for 15 min in PBS and incubated with secondary anti-rabbit Alexa Fluor 488-conjugated antibody at 1:500 for 1h at RT, protected from light. Following the washing step, tissues were counterstained with DAPI and mounted with anti-fade mounting medium (Sigma). Visualization was performed on Leica DM2000 fluorescent microscope

### Dynamic light scattering

In these studies, recombinant human vimentin was kindly provided from Josef Käs and Jörg Schnauss (University of Leipzig) and by Karen Ridge (Northwestern Univ). The pseudovirus size (hydrodynamic radius) was determined by dynamic light scattering (DLS) using a DynaPro 99 instrument (*77*). The method measures the diffusion constant of the particles using the autocorrelation function of fluctuations in scattered light intensity. The radius *R_h_* is calculated from the relation 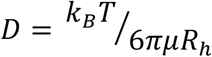, where *D* is the translational diffusion constant, *k_B_* is the Boltzmann constant, *T* is the temperature, and *μ* is the solvent viscosity. To examine the interaction between the SARS-CoV-2 pseudovirus and vimentin, we performed DLS measurements on pseudovirus solutions with and without addition of human recombinant vimentin. As a control for non-specific binding, in some cases, addition of DNA was used to determine how polyelectrolyte polymers other than vimentin interact with the pseudoviruses. Interaction of spike protein incorporated into the pseudovirus’s surface with vimentin oligomers would increase *R_h_* either by adding to the surface of a single pseudovirus or resulting in aggregation.

An estimate of the stoichiometry of vimentin oligomers to spike proteins is made from the concentration of pseudoviruses given by the manufacturer as 1.1×10^6^/mL, and the weight concentration and size of vimentin oligomers. Estimating the vimentin oligomer length as the hydrodynamic diameter, and the length of a vimentin unit length filament as 65 nm, leads to an estimate of 3 x 10^12^ vimentin oligomers/mL at the highest concentration. Assuming each pseudovirus contains 50 spike proteins leads to an estimate of 5 x 10^7^ spike proteins/mL. Therefore, at the highest concentration in the titration, there is a 6 x 10^4^x excess of vimentin oligomers to spike protein.

### Atomic force microscopy

A Multimode ^™^ AFM with a Nanoscope IIIa controller (Veeco, Santa Barbara, CA) operated in contact mode was used to image pseudoviruses before and after addition of vimentin or DNA. The cantilevers used were silicon nitride with a spring constant of 0.35 N/m (Veeco, Santa Barbara, CA). The AFM was calibrated using a 3D reference of 200 nm height and 10 μm pitch (Digital Instruments, Santa Barbara, CA). Deflection and height images were obtained with scan rates of 1 Hz with a resolution of 512 pixels/line. Briefly, for AFM evaluation of pseudoviruses in control (non-vimentin addition) and samples preincubated with human recombinant vimentin, a 10 μl drop of pseudoviruses or pseudoviruses/vimentin was applied on cleaved mica (SPI Supplies, West Chester, PA) and analyzed in a dried environment.

### Immunostaining of cell surface vimentin

To visualize surface vimentin, cells were fixed at approximately 50-70% confluency in 4% paraformaldehyde for 30 min at 37°C. Following fixation, to avoid non-specific binding cells were incubated with blocking-buffer (1% BSA in PBS) for 30 min at RT. After blocking, cells were incubated overnight at 4°C with primary rabbit monoclonal and polyclonal anti-vimentin antibodies at 1:500. Next, cells were incubated for 1h at RT with an Alexa Fluor 488-conjugated goat antirabbit secondary antibody at 1:1000. Counterstain was performed with DAPI nuclear stain. To differentiate surface vimentin from intracellular vimentin, additional staining was performed with addition of permeabilization step with 0.1% Triton X-100 in PBS for 15 minutes. Images were captured with a fixed-stage Zeiss Axio Examiner Z.1 microscope (Carl Zeiss Microscopy GmbH, Germany) and confocal scanning unit (Yokogawa CSU-X1, Yokogawa Electric Corporation, Japan).

### Antibodies

For fluorescent imaging of vimentin and for experiments using anti-vimentin antibodies to block host cell transduction by SARS-CoV-2 pseudovirus, we used the following anti-vimentin antibodies: (i) Pritumumab (Nascent Biotech), a human derived IgG antibody that targets the C-terminus of vimentin; (ii) Chicken polyclonal IgY antibody (Novus Biologicals, Cat# NB300-223); (iii) primary anti-vimentin monoclonal antibody developed in rabbit immunized with a 17-residue synthetic peptide from a region within human vimentin amino acids 425-466 (Abcam, Cat# ab92547); (iv) Rabbit Monoclonal (Cell signaling Technology, Cat#5471); and (v) Ser56 N-terminus aa 1-80 (primary anti-vimentin polyclonal rabbit antibody; antibodies-online.com, Cat# ABIN6280132).

### Pseudovirus description and manufacture

We used two independent sources of pseudovirions. The first were commercially available SARS-CoV-2 GFP-reporter pseudovirus from Integral Molecular (Integral Cat# RVP-701G). These pseudoviruses display antigenically correct spike protein pseudotyped on replication-incompetent virus particles that contain a heterologous lentiviral (HIV) core. They were produced in HEK-293T cells using three separate plasmids, encoding the spike protein, a lentiviral gag polyprotein, and a reporter gene.

The second source was from in-house manufacturing performed in the Pyrc laboratory. The pseudoviruses were produced as previously described (*78*). Briefly, HEK293T cells were seeded on 10 cm^2^ cell culture dishes, cultured for 24 hr at 37°C with 5% CO2 and transfected using polyethyleneimine (Sigma-Aldrich, Poland) with the lentiviral packaging plasmid (psPAX), the VSV-G envelope plasmid (pMD2G) or SARS-CoV-2 S glycoprotein (pCAGGS-SARS-CoV-2-S) and third plasmid encoding GFP protein (Lego-G2). Cells were further cultured for 72 hr at 37°C with 5% CO_2_, and pseudoviruses were collected after 48 and 72 h, further filtered by 0.22 μm filter, and 10-times concentrated with Amicon Ultra-15 Centrifugal Filters (10k, Merck Millipore) and stored at −80 °C.

### In vitro transduction of cells by pseudoviruses

A549-ACE2 and HEK 293T-ACE2 cell lines were used as infection hosts for SARS-CoV-2 GFP-reporter pseudovirus (Integral Molecular, Integral Cat# RVP-701G). To transduce cells with the pseudovirus, cells were seeded in 96-well plates at approximately 50% confluency and pseudoviruses were introduced to cells by pipetting 50 μL (at 10^9^ pseudoviruses/mL) into each well and in some cases by a serial 1/2 dilution of the pseudovirus. To assess the involvement of vimentin during SARS-CoV-2 infection, anti-vimentin antibodies were introduced into the wells either at the same time as the pseudovirus or for a short incubation time prior to pseudovirus exposure. The pseudoviruses were either washed out after a 2 hr exposure time interval or were allowed to incubate with the cells throughout the remainder of the experiment. Cells were incubated for 72h at 37°C with 5% CO2.

### Pseudovirus entry characterization

Pseudovirus entry was measured in one of two ways: the first is via epi-fluorescence microscopy and the second is by a microplate reader. In the first case, the percentage of cells infected by SARS-CoV-2 pseudovirus was quantified by calculating the ratio of cells expressing GFP to the total number of cells. Cells were imaged using a Nikon Eclipse Ti (Nikon Instruments) inverted microscope equipped with an Andor Technologies iXon em+ EMCCD camera (Andor Technologies). Cells were maintained at 37°C and 5% CO2 using a Tokai Hit (Tokai-Hit) stage top incubator and imaged using a 10x objective Pan Fluor NA 0.3 (Nikon Instruments) at 24 hr, 48 hr, and 72 hr time points after pseudovirus transductions. In some cases, nuclei were stained using Hoechst prior to imaging to facilitate counting total cell numbers. The total number of cells and the number of cells expressing GFP were manually counted in randomly selected 65 μm by 65 μm imaging windows in ImageJ. The percentage of cells infected is calculated by taking the ratio of the number of cells expressing GFP to the total number of cells. Results were normalized to untreated control samples.

For studies using a microplate reader, A549-hsACE2 and HEK293T-hsACE2 cells were seeded in 96-wells optic-bottom black cell culture plates, maintained for 24 h at 37°C with 5% CO2 and transduced with pseudoviruses harboring VSV-G or S-SARS-CoV-2 proteins or lacking the envelopeprotein (ΔEnv) in the presence of polybrene (4 μg/ml; Sigma-Aldrich, Poland), and rabbit anti-vimentin antibody at 1:1000, 1:500 and 1:100 dilutions (Abcam, Cat# ab92547) or control PBS. After 4 hr incubation at 37°C unbound virions were removed by washing thrice with PBS, and cells were further cultured for 72 h at 37°C with 5% CO2. After 72 hr supernatant was removed and cells were washed carefully with PBS. 100 μl of PBS was added to each well and fluorescence was measured with Varioskan LuX microplate reader (ex= 480, em= 530). Control fluorescence values in Spike-GFP and VSV-G-GFP were normalized to 1. Results were normalized to untreated control samples.

### Endocytosis model

To examine the interaction between extracellular vimentin and the SARS-CoV-2 virus, we developed a coarse-grained molecular dynamics-based 3D model, modeling the role of extracellular vimentin as an attachment factor. The model consists of the cell membrane, extracellular vimentin, the angiotensin-converting enzyme 2 (ACE2) receptor, the SARS-CoV-2 virus, and the interactions between them. A portion of the cell membrane (approximately 500 nm x 500 nm in area) is modeled as a tethered membrane with a bending rigidity of 40k_B_T. Based on prior experiments (*48*), the extracellular vimentin is considered to be a 40 nm semiflexible filament, bound to the surface of the cell membrane. For simplicity, we have assumed that the SARS2 virus is fully bound to the ACE2 receptor: the spike-ACE2 interaction is given a stiff harmonic spring as ACE2 has a high affinity towards spike protein (*1*). We designed the SARS-CoV-2 virus as a 100 nm diameter elastic viral spherical shell, and viral shell is decorated with 200 spikes (an upper bound estimate), each containing sticky sites that bind to extracellular vimentin through an attractive Lennard-Jones potential. For excluded volume interactions, we implement soft-core repulsion springs should any two particles begin to overlap. Each simulation is run for a total of 10^8^ simulation units corresponding to 50 s. We defined a degree of membrane wrapping metric to reflect membrane curvature as a measure of the depth of the bowl generated by the cell membrane. Specifically, we measure the distance between the height of the cell membrane (far from the bowl) versus the height of the deepest invagination of the membrane bowl: this value is then normalized by the total height of the SARS-CoV-2 virus, 120 nm. (See SI Materials for full details on the model.)

### Acquisition of vimentin released from neutrophils

To assess whether vimentin may be acquired from extracellular environment, supernatant from phorbol myristate acetate (PMA) stimulated neutrophils were added to the vim-null mEF. To do so, blood was obtained from healthy volunteers (protocol number for human blood draw: R-I-002/231/2019) and neutrophils were isolated by gradient density separation with use of Polymorphprep. Isolated neutrophils were counted and transferred to Eppendorf tubes containing DMEM at concentration 6×10^5^ cells/mL. 100 nM of PMA was added to neutrophils for 30 min at 37°C. After incubation, cells were centrifuged and 100 μl of supernatant was added to a prewashed well with 1.5×10^4^ vim-null mEFs in a black 96-well glass bottom plate for 30 minutes. In the next step, cells were washed three times, fixed and immunofluorescently stained as described above.

### Statistical methods

Data are presented as mean values ± SEM unless otherwise stated. Each experiment was performed a minimum of two times unless otherwise stated. The unpaired, two-tailed Student’s t test at the 95% confidence interval was used to determine statistical significance. Denotations: *, P ≤ 0.05; **, P < 0.01; ***, P < 0.001; NS, P > 0.05.

## Acknowledgements

We acknowledge Robert Goldman, Karen Ridge, and Nav Singh for insightful discussions. This work was supported by NSF MCB 2032861 award to AP and JMS., NIH GM136259 award to PAJ., and the National Science Center of Poland: UMO-2020/01/0/NZ6/00082 awarded to RB. DG acknowledges funding from Syracuse University SOURCE grant.

## Author contributions

LS, MS, RB and AP performed and analyzed the SARS-CoV-2 pseudovirus transduction studies. ML and KP designed and manufactured pseudovirions. DG and DI contributed to pseudovirus transduction analysis. LS, MS, JR, NM, and AP performed extracellular vimentin staining. RB, PJ, and RC performed and analyzed dynamic light scattering experiments. FB performed and analyzed the atomic force microscopy measurements. SG and JMS developed the endocytosis model and analysis. KP, PJ, JMS, RB, MS, JR and AP wrote the manuscript. RB and AP initiated and oversaw the entire project.

## Supplementary Figures

**SI Fig. 1.**
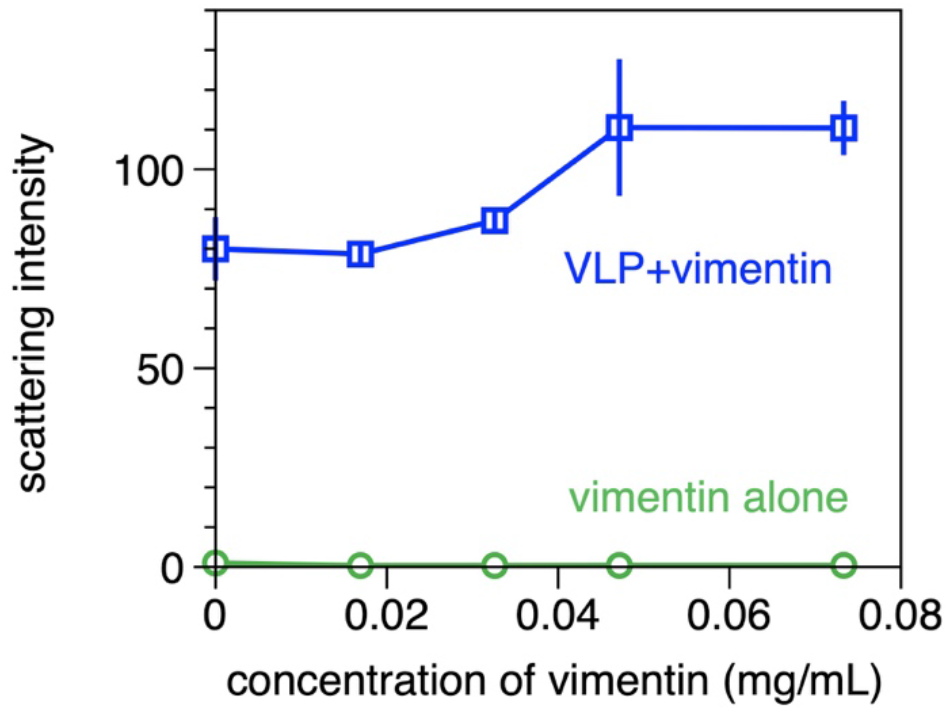
The scattering intensity of the viral parties was much larger than that of the added vimentin and was increased after adding vimentin.

**SI Fig. 2.**
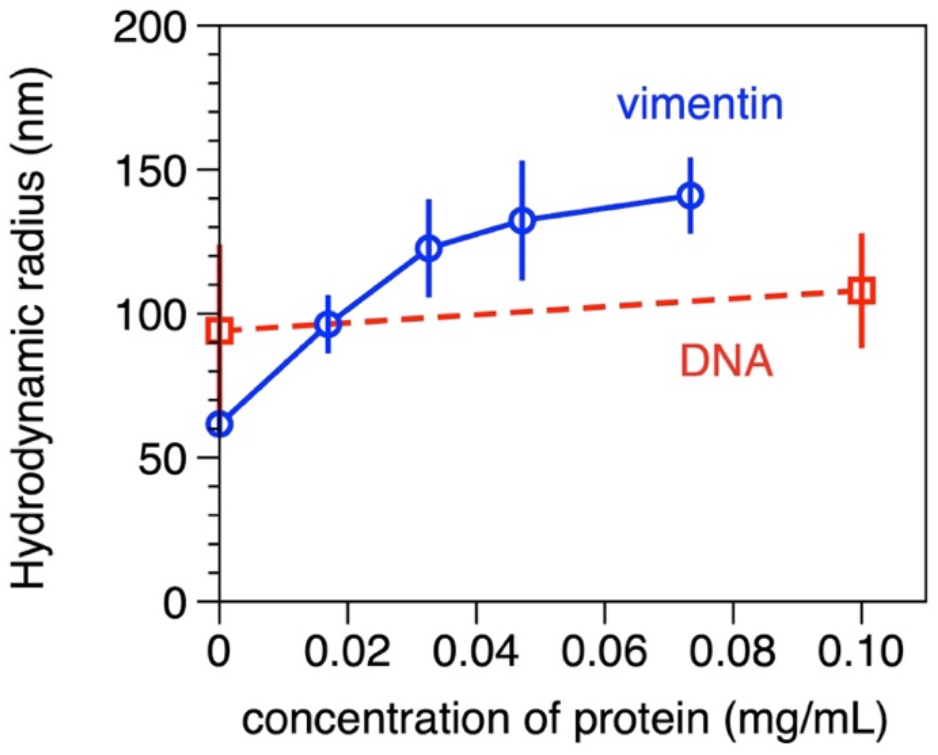
Dynamic light scattering data showing the relative increase of pseudovirion particles with addition of either vimentin or DNA. Addition of vimentin increases relative size of the particles whereas DNA does not.

**SI Fig. 4.**
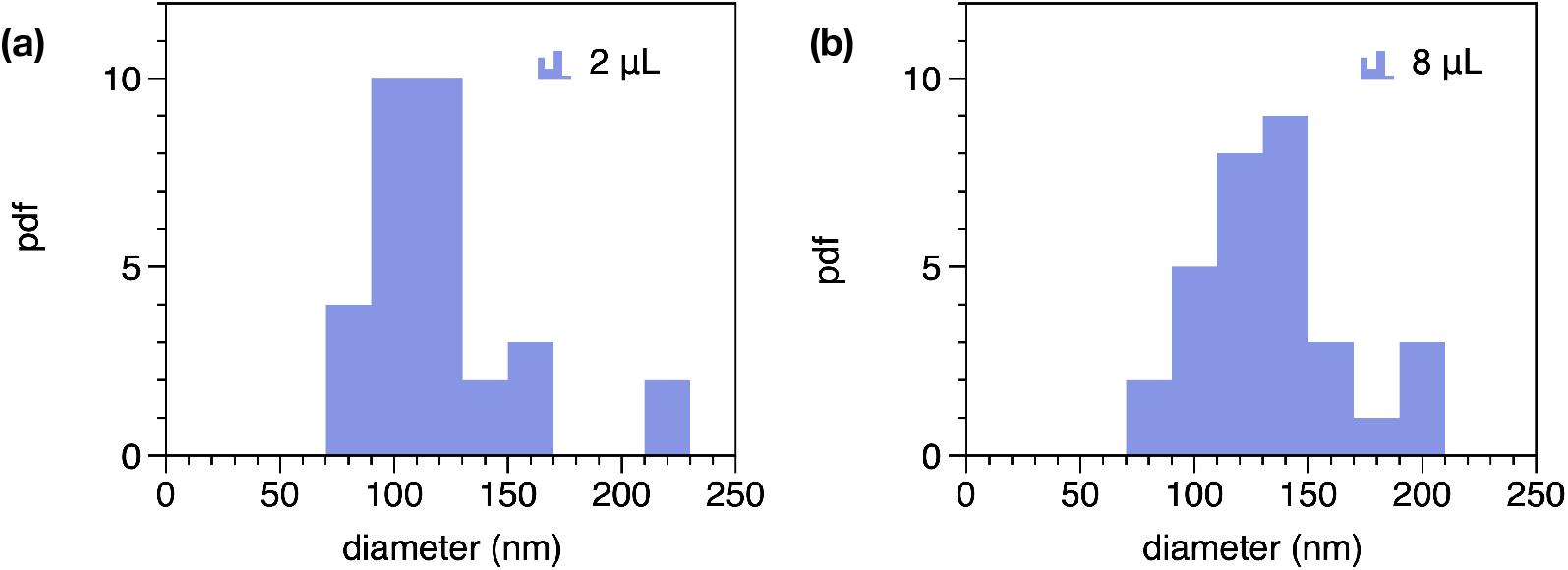
The probability distribution function of particle diameter for SARS-CoV-2 pseudovirus samples after the addition of either (a) 2 μL or (b) 8 μL of 10 mg/mL DNA. Particle sizes were measured by atomic force microscopy.

**SI Fig. 5.**
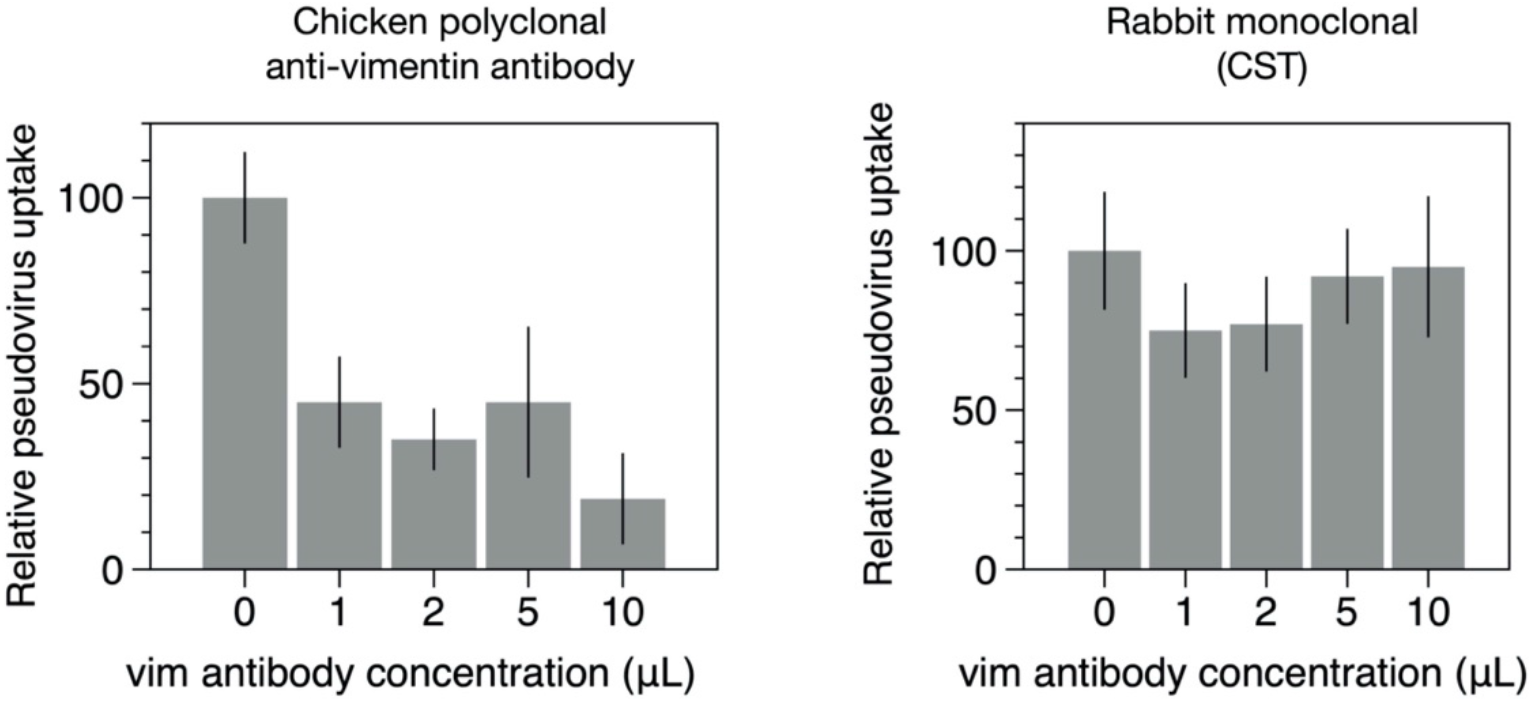
Results from the anti-vimentin rabbit monoclonal antibody from Cell Signaling Technologies does not block uptakes of SARS-CoV-2 pseudovirus in HEK 293T-hsACE2 cells. Error bars represent mean ± standard deviation.

**SI Fig. 6.**
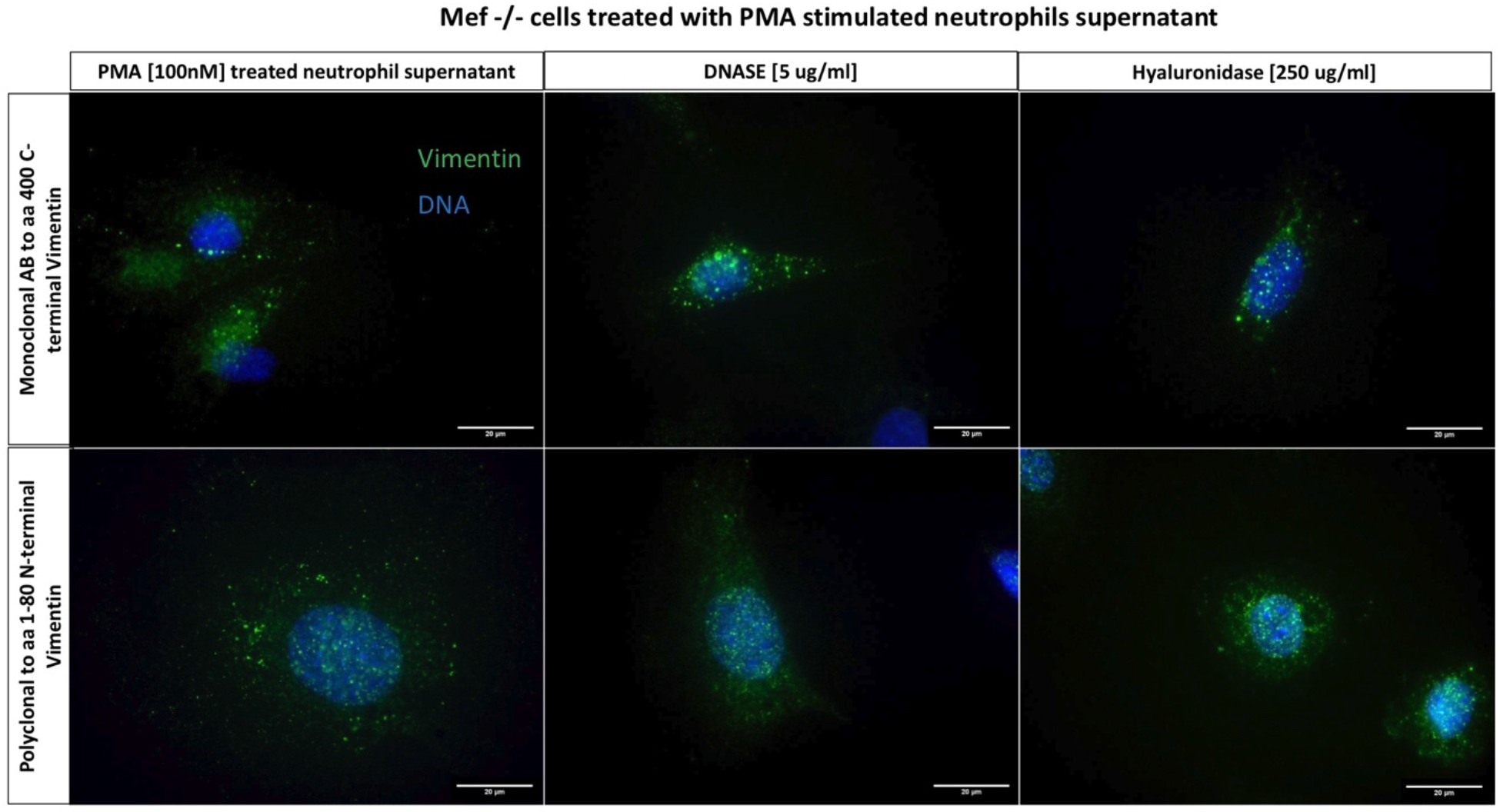
Immunofluorescence images indicating presence of cell surface vimentin on vim -/- mEF after exposure to PMA-stimulated neutrophil supernatant. Neither DNASE or hyaluronidase treatment alters the presence of cell surface vimentin.

## Supplemental Information

### A. Modeling cell surface vimentin, SARS-CoV-2 and cell-membrane interactions

To test the notion that extracellular vimentin bound to the cell membrane can initiate endocytosis of the SARS2 virus, we developed a multiscale, coarse-grained molecular dynamics-based computational model. The model consists of a cell membrane, extracellular vimentin (ECV), the angiotensin-converting enzyme 2 (ACE2), and a virus containing spike proteins.

The cell membrane is modeled as a self-avoiding, tethered sheet with fixed connectivity as shown in Fig. [1]. It is a network of equilateral triangles consisting of particles as *N* nodes and tethers to connect them. The two-dimensional sheet is embedded in three-dimensional Euclidean space. We introduce self-avoidance to the system by satisfying the condition 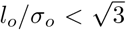, where *l_o_* is the edge length of a triangle in the mesh and *σ_o_* is the diameter of a particle [1]. Stretchability of the membrane is encoded in nearest-neighbor harmonic springs with the energy

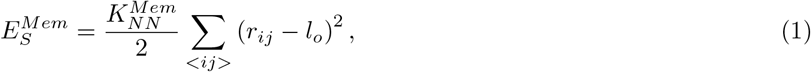

with spring constant 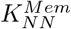 and < *ij* > represents all nearest-neighbor nodes *i* and *j*. We also have an explicit bending rigidity modeled by adding another harmonic spring between every second nearest neighbor of each node [2, 3] having energy

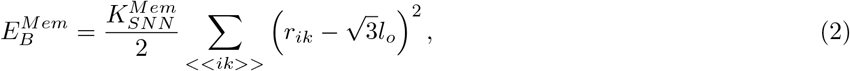

with ≪ *ik* ≫ denoting all second-nearest neighbors. Spring constant 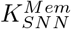 can be converted to a bending rigidity

Given the experimental findings of non-filamentous extracellular vimentin bound to the cell membrane, we focus on extracellular vimentin tetramers [4]. The extracellular vimentin tetramers are represented as semiflexible filaments, containing both springs between two consecutive monomers with spring constant, 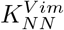, and angular springs between three consecutive monomers to capture the bending rigidity with angular spring constant, 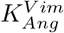. The energy for each monomer triplet is, therefore,

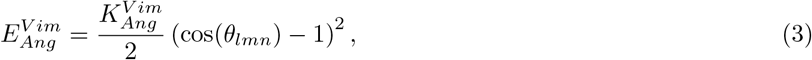

where *l, m, n* denote three consecutive monomers along the filament. For simplicity, we have assumed the ACE2 receptor is the same length as the vimentin tetramers and is also a semiflexible filament with corresponding two-body springs and angular springs, 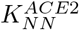 and 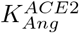, respectively. The ACE2 receptor is stiffer than vimentin and provides stability during the wrapping process. Both the vimentin tetramers and the ACE2 receptor are also connected with the membrane via springs as shown in Fig. [2] with the same two-body spring constant, which gives them the freedom to bend at any angle with respect to the membrane. We place extracellular vimentin randomly on the membrane with coverage *ϕ_Vim_*, whereas ACE2 is located at the center of the membrane. The extracellular vimentin now becomes cell-surface vimentin.

The virus is modeled as a deformable shell with spikes. The shell is constructed as a Fibonacci sphere, where particles/monomers are placed on the sphere in a spiral. We use a Delaunay triangulation to find triangles amongst the particles and their corresponding edges. These particles are also connected via nearest-neighbor harmonic springs each with spring constant, 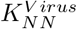 to arrive at a tethered spherical membrane with fixed connectivity. Spike proteins are homogenously placed on the shell surface as shown in Fig. [3]. The spike protein filaments also have harmonic spring potential between two consecutive monomers with spring constant 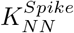. Their bending rigidity is also coded via 3-body, or angular, spring with spring constant, 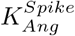. The spike protein filaments can take any angle with respect to the virus [5, 6].

To take into account excluded volume interactions, we implement a soft-core repulsion spring between all monomers with no other interactions via energy

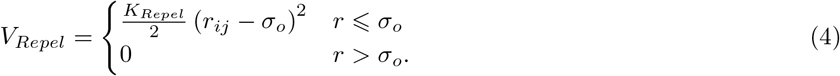

**FIG. 1.**
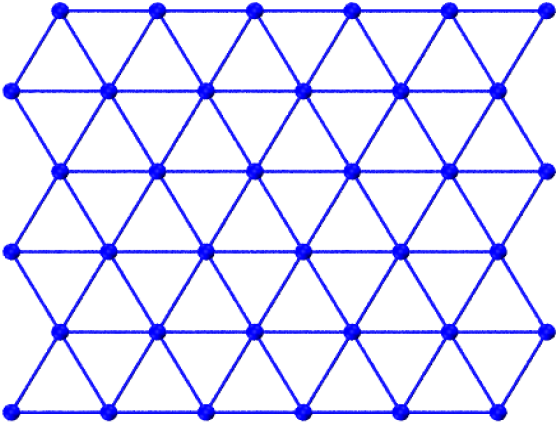
*Tethered membrane as a cell membrane*. Blue particles denote the nodes of the triagular mesh and blue lines denote the tethers.

**FIG. 2.**
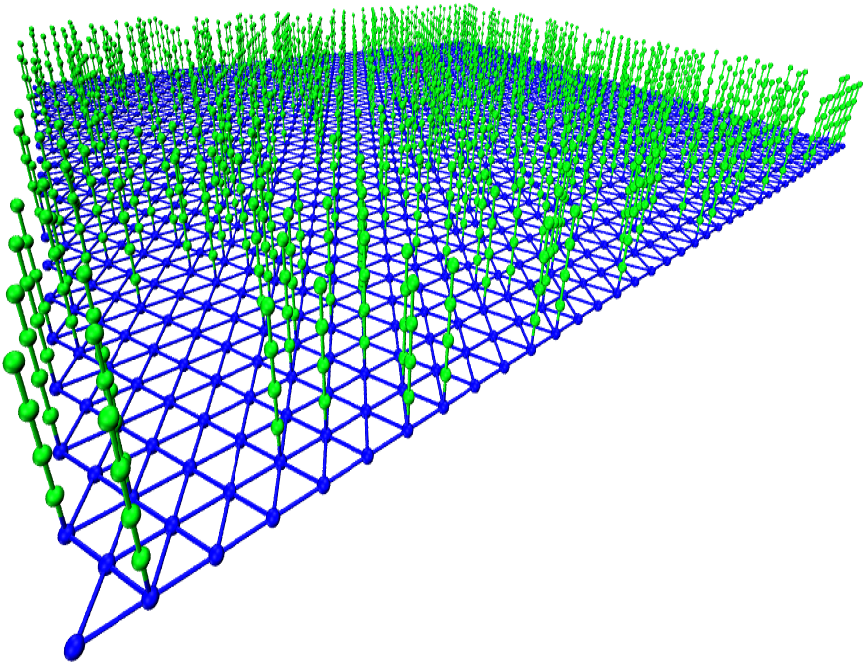
*Cell membrane with bound extracellular vimentin*. Extracellular vimentin (ECV) tetramers are denoted in green. ECV tetramers are randomly attached to the cell surface to become cell surface vimentin. The coverage of extracellular vimentin depicted here is *ϕ_Vim_* = 0.4, i.e., 40 percent of the membrane nodes have bound ECV.

As for other interactions, the spike-ACE2 interaction is modeled as a stiff harmonic spring between the two filament ends with spring constant 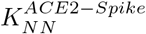 since ACE2 has a high affinity to the spike protein [7, 8]. Additionally, there is an attractive Lennard-Jones potential between the spike protein and the cell surface vimentin with a higher cut-off at 2*σ_o_* as given by

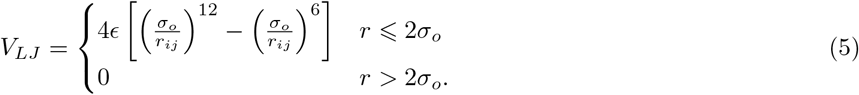

To convert our simulation units to biological units, we use 1 simulation unit length = 10 *nm*, 1 unit simulation time= 10^−3^ *s* and 1 unit force = 10 ^−1^ *pN*. From these scales, we can define all the parameters in biological units. The diameter of the virus is 100 nm, which is very similar to our own DLS measurements and similar to SARS2 [9–11]. The size of the virus is small compared to the size of the cell, typically ~ 20 *μm* in diameter, which is about 200 times bigger than the virus. Thus, during endocytosis, the virus is interacting with a small patch of the cell membrane. Hence, we simulated a tethered sheet of *length* × *width* = 55 nm × 480.6 nm, which is a small surface area of the cell membrane. This cell surface is covered with cell surface vimentin tetramers of length 40 *nm*, which is slightly smaller than the more typical 60 — 90 nm. We vary the density of the extracellular vimentin (ECV) to quantify its effect on viral uptake. The number of spike proteins on the viral surface can be different based on virus size [12]. We consider 200 spikes[9, 11, 13, 14] on the virus each of length 20nm [5, 9, 15] and diameter 10nm [11], to which ECV can attach. The bending rigidity of cell membrane is approximately 40kB*T* based on the scales we have defined. Finally, each simulation was run for 10^8^ simulation time units, which corresponds to 50 *s* with recording trajectory data every 25 ms. See Table 1 for the parameters used in the simulations.

**FIG. 3.**
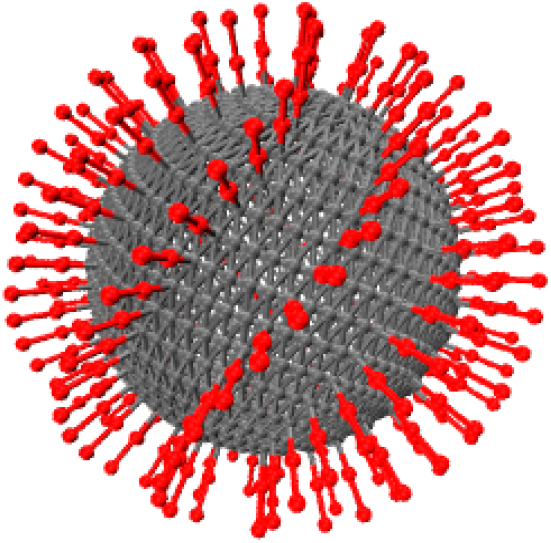
*SARS2 virus*. Spike proteins are placed homogeneously on the surface of the virus.

For each set of parameters, 10 realizations were computed and used to compute average quantities of the fraction of spike proteins bound to ECV, and the degree of wrapping by the cell membrane, as defined in the Methods section of the manuscript. All computed quantities indicate the extent of endocytosis, or wrapping, of the virus by ECV.

**TABLE I.** Table of parameters used unless otherwise specified.

| Parameters | Value | Reference |
| --- | --- | --- |
| $K_{NN}^{Mem}$ | $1 pN/nm$ | [16] [17] |
| $K_{SNN}^{Mem}$ | $0.1 pN/nm$ | [18] [19] |
| $K_{NN}^{Virus}$ | $50 pN/nm$ | SARS Cov-2[20],<br>Brone mosaic virus<br>[21], influenza<br>virus[22], Human<br>Adenovirus [23]<br>Deformable NP[24] |
| $K_{NN}^{Spike}$ | $0.1 pN/nm$ | [25] |
| $K_{NN}^{Vim}$ | $1 pN/nm$ | [26] |
| $K_{NN}^{ACE2}$ | $5 pN/nm$ | - |
| $K_{NN}^{ACE2-Spike}$ | $5 pN/nm$ | [27] |
| $K_{Ang}^{ACE2}$ | $500 k_B T$ | - |
| $K_{Ang}^{Vim}$ | $100 k_B T$ | - |
| $K_{Ang}^{Spike}$ | $100 k_B T$ | - |
| $\epsilon$ | $10 k_B T$ | - |
| $K_{Repel}$ | $1 pN/nm$ | - |

